# The effects of different frequencies of rhythmic acoustic stimulation on gait kinematics and trunk sway in healthy elderly population

**DOI:** 10.1101/2020.11.20.390955

**Authors:** Roberta Minino, Emahnuel Troisi Lopez, Pierpaolo Sorrentino, Rosaria Rucco, Anna Lardone, Matteo Pesoli, Domenico Tafuri, Laura Mandolesi, Giuseppe Sorrentino, Marianna Liparoti

## Abstract

The use of rhythmic acoustic stimulation (RAS) in improving gait and balance in healthy elderly subjects has been widely investigated. However, methodologies and results are often controversial. In this study, we hypothesize that both the kinematic features of gait and stability, depend on the frequency at which RAS is administered. Our aim was to observe, through 3D Gait Analysis, the effect of different types of RAS (at a fixed frequency or based on the average cadence of each subject) on both gait spatio-temporal parameters and stability. The latter was estimated through an innovative measure, the trunk displacement index (TDI) that we have recently implemented. We observed that the low frequencies RAS led to a general slowdown of gait, which did not provide any clear benefit and produced harmful effects on stability when the frequency became too low compared to the individual natural frequency. On the contrary, the high frequencies of RAS showed a slight acceleration of gait, accompanied by better stability (as documented by a lower TDI value), regardless of the type of RAS. Finally, the RAS equal to the individual natural cadence also produced an increase in stability.

## 1. INTRODUCTION

Aging-related motor impairment arises from multiple factors, including bone loss, muscle atrophy, and a decline of both central and peripheral nervous systems functioning^1^. As a consequence, the elderly people show a reduction of the static and dynamic balance, slower movements, altered modulation of strength and increased walking variability^2–4^. All these factors are closely related and contribute to increased instability and risk of falls^5^. Due to the frailty of elderly people, falling carries a high burden, as it compromises personal autonomy, limits mobility and increases morbidity and mortality. Indeed, several studies highlighted the close connection and the increasing rate of death originating from fall events, addressing it as an important public health problem^6–8^. Additionally, in the elder population, cognitive decline and some psychological factors, such as stress, anxiety and depression contribute to an increased risk of falling and a worsened quality of life^9^. Hence, finding preventive strategies is needed.

In order to quantify instability, identify weak points and try to minimize these risks, several studies have investigated movement and, in particular, locomotion in elderly subjects, using different methodologies and tools^10–12^. Among these, the 3D gait analysis (3D-GA), typically considered the gold standard^31^, is a quantitative method to measure the kinematics of movements, widely applied for the assessment of motor skills, in physiological and pathological conditions^13–19^.

Many studies have investigated the strategies adopted to counterbalance the increased (static and dynamic) instability, in physiological or pathological conditions. Among these, several works focused on the effects of sensory stimulation on balance and locomotion^20–23^. Sensory afferents provide information on the reciprocal position and movement of body segments and their orientation in space^20^. The increase in sensory information (auditory, visual or vibro-tactile stimulations) can help static and dynamic balance, even for individuals with sensory and motor impairment^24^. Accordingly, it has been widely demonstrated that an appropriate acoustic stimulus improves walking parameters in different pathological conditions, including Parkinson’s disease^25–27^, stroke and multiple sclerosis^28^.

Given the therapeutic potential, some effort has been devoted to the description of the mechanisms through which acoustic stimuli, and rhythmic acoustic stimulation (RAS) in particular, influence walking. This is particularly relevant for the possibility of using a device based on RAS to support motor rehabilitation even in elderly people with a high risk of falls. Dickstein et al. studied how walking synchronizes to an acoustic stimulus emitted by a metronome, at different fixed (i.e. irrespective of each participant’s baseline frequency) frequencies (60, 110, and 150 beats/min). The authors showed that lower frequencies display greater ease of synchronization^29^. Conversely, Yu et al. explored the effects of three different frequencies based on the cadence of each participant (i.e. 90%, 100% and 110% of the average cadence of each participant), on spatiotemporal gait parameters^30^. It was suggested that the faster stimulus affects the most gait parameters, such as the stride length, the cadence and the walking speed, as compared to the non-cued walking. Hence, how to most effectively set the stimulus features remains an open question, as it is not clear which gait parameters and cortical processes are involved in the adaptation to acoustic stimulation.

Most of the studies have focused selectively on the lower limbs, excluding the other body segments^14^. However, in order to maintain balance^32^, walking entails through synergistic and coordinated movements of the upper limbs, trunk and lower limbs^33^. Importantly, central balance control does not intervene on each muscle individually, but rather aims at the control of the centre of mass (COM), with segmental or sub-segmental adjustments left to hierarchically lower mechanisms^32,34,35^. Therefore, studying the static and dynamic balance of a population with higher risk of falling, such as the elderly, through the exclusive study of the lower limbs, is highly limiting.

We have recently implemented an innovative measure of postural stability which assesses the trunk displacement in relation to the centre of mass, that we named trunk displacement index (TDI). This measure, obtained from 3D-GA data, evaluates the ratio of the displacements of trunk and COM on three anatomical planes, conveying the overall motor performance. Considering that a less efficient control of the COM is related to increased risk of falling, we hypothesize that the effectiveness of different acoustic stimuli on the gait stability can be evaluated assessing the TDI value.

The aim of this work is to evaluate the effect of different frequencies (fixed or established on the average cadence of each subject) of RAS on walking in healthy elderly subjects. As shown, studies on the effect of RAS on gait used a variety of methods producing conflicting results. We hypothesized that optimal gait stabilization does not occur at the same frequency for each subject, rather, the optimal frequency of stimulation is subject-specific and span around the individual cadence of each subject. In order to test our hypothesis, we used 3D-GA to calculate spatio-temporal gait parameters and TDI in three experimental conditions characterized by a RAS frequency lower, equal or higher than that of each subject^34^. More precisely, each subject was recorded while walking at a RAS frequency which was: 1) equal to his/her cadence, 2) lower (90% of the basal cadence and 80 bpm) and 3) higher (110% of the basal cadence and 120 bpm). Furthermore, we carried out the correlation analysis in order to evaluate the relationship between the gait parameters and stability, expressed as TDI.

## 2. METHODS

### 2.1 Participants

Twenty-two elderly people were recruited (table 1), nine of which were excluded because they did not meet the following inclusion criteria: *(i)* 65 to 85 years old; *(ii)* no skeletal, muscular and neurodegenerative disorders; *(iii)* no hearing impairment; *(iv)* Beck Depression Inventory (BDI)^36^ score < 13; *(v)* Mini Mental State Examination (MMSE) score > 23,80^37^; *(vi)* Frontal Assessment Battery (FAB) score > 12.03^38^. Thirteen participants (6 males and 7 females) were included. Written informed consent was obtained from all participants prior to participation, in accordance with the declaration of Helsinki. The study was approved by the local Ethic Committee (University of Naples Federico II, n. 26/2020).

**Table 1.** Demographic, anthropometric and neuropsychological participant’ characteristics (mean ± standard deviation). Weights in kilograms, heights in centimetres, body mass index (BMI), mini mental state examination (MMSE), frontal assessment battery (FAB), Beck’s depression inventory (BDI).

| Participant Characteristics |  |
| --- | --- |
| Demographic Data |  |
| Ages (years) | 73.4 (±5.7) |
| Education (years) | 11.6 (±4.8) |
| Gender (m/f Ratio) | 6/7 |
| Anthropometric Parameters |  |
| Weights | 65.27 (±7.57) |
| Heights | 161.78 (±8.89) |
| BMI | 25.0 (±2.8) |
| Neuropsychological Evaluation |  |
| MMSE (adjusted) | 27.65 (±1.85) |
| FAB | 16.41 (±1.66) |
| BDI | 6.85 (±3.98) |

### 2.2 Gait Analysis Assessment

#### 2.2.1 Protocol

The 3D-GA was carried out in the Motion Analysis Laboratory of the University of Naples “Parthenope”. The 3D-GA data were acquired through a Stereophotogrammetric system, equipped with eight infrared cameras (ProReflex Unit-Qualisys Inc., Gothenburg, Sweden). In agreement with the modified Davis protocol^39^, fifty-five passive markers were positioned on each participant on anatomical landmarks of the feet, the lower limb joints, the pelvis and the trunk, as well as on the upper limb joints and on the head (Fig.1). The recorded data were processed using a tracking data software (Qualysis Track Manager by Qualisys AB, Göteborg, Sweden) and a software (Visual 3D by C-Motion Inc., Germantown, MD) to rebuild a model of the skeleton^16,40^. We recorded four trials for each frequency and each trial included four consecutive steps (a gait cycle), similarly to Liparoti et al.,^15^. In order to obtain a more reliable estimate, we calculated the average of each step within a trial. Participants were asked to walk at the pace of the acoustic stimulus, emitted by a metronome (MA-1 Solo Metronome, Korg -UK). The acoustic stimulation was set to have a tone of 440 Hz^41^(such as a metronome tic), with an easily audible volume^42^.

**Fig. 1.**
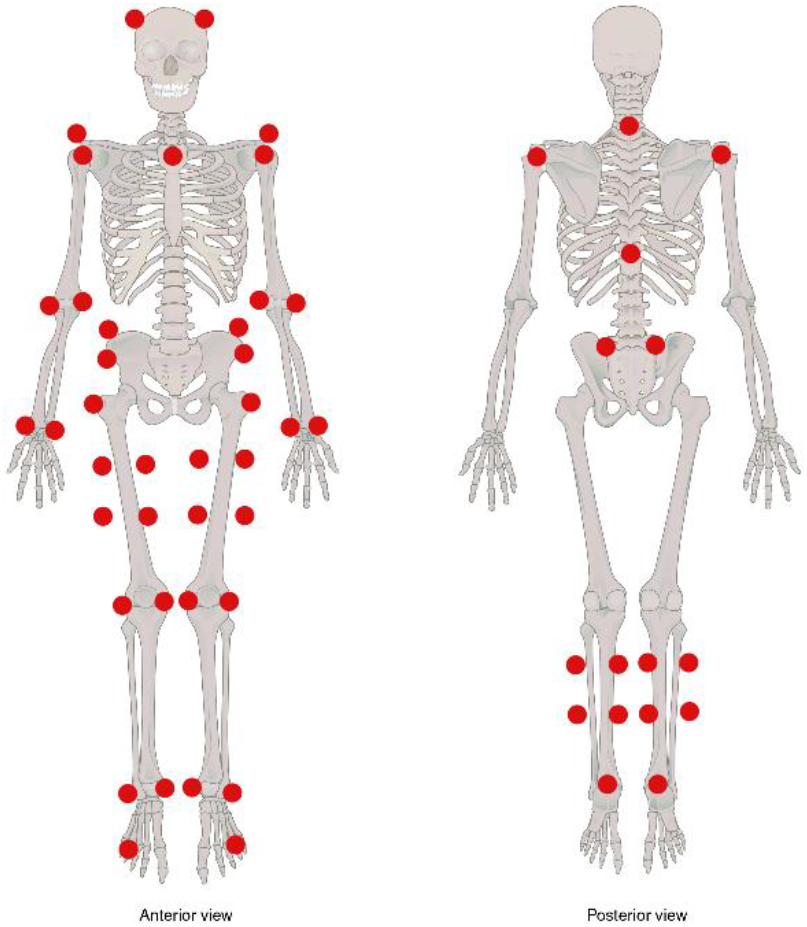
Marker set of the fifty-five markers positioned on the anatomical landmarks of the feet, lower limb joints, pelvis, trunk, upper limb joints and on the head. Adapted from “Axial and Appendicular Skeleton” (http://cnx.org/content/col11496/1.6/) by OpenStax College, used under CC by 4.0 (https://creativecommons.org/licenses/by/4.0/)

The data acquisition was carried out in two times. During the first one, subjects were recorded at a self-selected cadence in order to calculate the natural cadence of each participant. In the second phase, we recorded the participants’ walking in 6 experimental conditions: 1) simple walking (SW); 2) walking with RAS at frequency equal to the subject-specific average cadence (AC) (100% AC); 3) walking with RAS at a frequency equal to 90% of the participants’ cadence (90% AC); 4) walking with RAS at a frequency equal to the 110% of the participants’ cadence (110% AC); 5) walking with RAS at fixed frequency equal to 80 beat per minute (bpm); 6) walking with RAS at fixed frequency of 120 bpm (table 2). The order of the acoustic stimuli was randomized to reduce the learning effect and fatigue. For the SW, all participants were instructed to walk at a normal pace over a 10-meter-long carpet.

**Table 2.** Frequencies based on average cadence of each participant measured in beats per minute (mean ± standard deviation). The 100%AC is the frequency equal to the average cadence of each participant. The 90% AC is the frequency equal to 90% of the participants’ cadence. The 110% AC is the frequency equal to 110% of the participants’ cadence.

| Participants | 100% AC<br>frequency<br>(bpm) | 90% AC<br>Frequency<br>(bpm) | 110% AC<br>Frequency<br>(bpm) |
| --- | --- | --- | --- |
| 1 | 115 | 103 | 126 |
| 2 | 117 | 104 | 128 |
| 3 | 106 | 95 | 117 |
| 4 | 113 | 102 | 124 |
| 5 | 105 | 95 | 115 |
| 6 | 90 | 81 | 99 |
| 7 | 109 | 98 | 120 |
| 8 | 129 | 116 | 142 |
| 9 | 110 | 99 | 121 |
| 10 | 100 | 90 | 110 |
| 11 | 108 | 97 | 119 |
| 12 | 119 | 107 | 131 |
| 13 | 112 | 100 | 123 |
|  | 110.23<br>(±9.46) | 99<br>(±8.40) | 121.15<br>(±10.37) |

#### 2.2.2 Spatio-temporal Parameters

3D-GA has been used to obtain the temporal and spatial parameters of gait. The former included Speed, Stance Time, Swing Time, Cycle time and Double Limb Support time (DLS). The latter included Stride Width and Stride Length. The variability coefficient (CV) (the ratio between standard deviation and average value x 100 (%)) was calculated for all spatio-temporal parameters, except speed.

#### 2.2.3 Trunk Displacement Index

In order to evaluate the stability, we calculated the TDI^34^, a newly described synthetic index,. We measured the trajectory of the centre of mass (*COMt*) in the three-dimensional space, during gait and calculated its mean 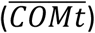. Thereafter, we calculated the same three-dimensional trajectory for the upper trunk (*Tt*). Subsequently, we calculated separately the distances between *COMt* and *Tt* from 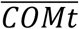 in each registered frame of the gait cycle of the individual, obtaining a vector of distances (*COMd* and *Td* respectively) for each one of the three planes, as shown in equations (1 and 2):

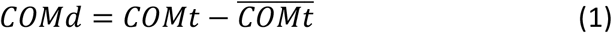

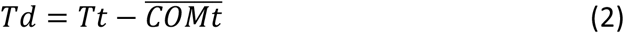

Finally, to enclose all the information in a unique parameter, which conveys the three-dimensional displacement of the trunk, we performed the following two final steps. Firstly, we summed the norm of each vector of distances (*COMd* and *Td*). Then, we calculated the ratio between those two values obtaining a unique dimensionless number, as shown in the equation (3), representative of the relationship between the three-dimensional displacement of the trunk and the COM^34^.

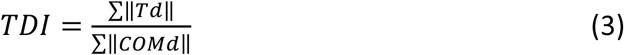

### 2.3 Statistical analysis

The statistical analysis was performed in Matlab (Mathworks^®^, version R2018b). The means and standard deviations of the gait parameters were calculated after correcting each variable by the body mass index (BMI) of each subject. Gait parameters were compared among all conditions, analysing three subsets separately: the first subset included the SW group and the high frequency stimuli groups (110% AC e 120 bpm); the second subset included the SW group and the low frequency stimuli groups (90% e 80 bpm); the third subset included the SW and the 100% AC groups.

The Shapiro-Wilk test was used to check the normal distribution of variables. Given the heterogeneous distribution of the variables and the small size of the sample, we performed nonparametric statistical testing. Friedman test was used to investigate the differences within subgroups, while Wilcoxon signed rank test was used to perform the pairwise comparison. Spearman correlation tests were carried out to test the relationship between TDI and gait parameters including all conditions.

## 3. RESULTS

### 3.1 Simple Walking – 100% AC frequency

Regarding the Temporal Parameters, there was a statistically significant difference in the DLS time (p < 0.01) between SW and 100% AC, with the latter showing higher values. About the Spatial Parameters, at 100% AC it was observed increased Stride Length (p < 0.05), as well as reduced DLS CV (p < 0.01), as compared to SW. Moreover, at the 100% AC the TDI was decreased as compared to SW (p < 0.01) (Fig. 2).

**Fig. 2.**
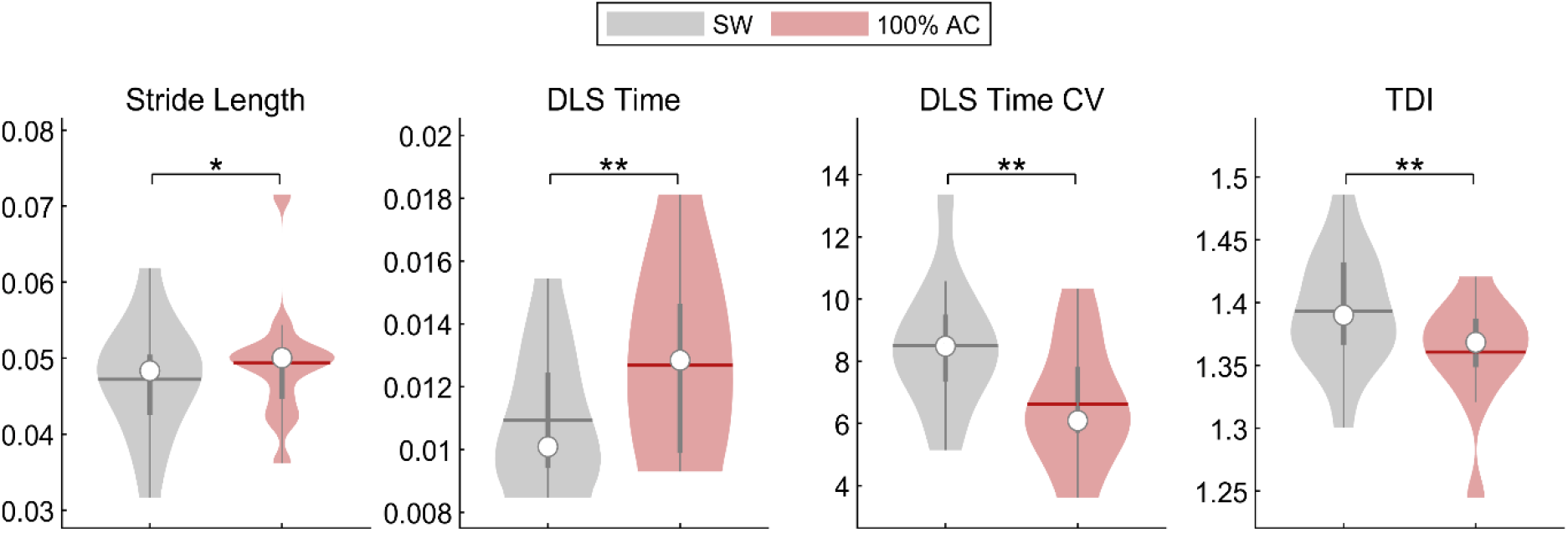
Spatio-temporal analysis of gait and Trunk Displacement Index (TDI). SW vs. 100% AC. Violin plots of spatio-temporal parameters and TDI value. Comparison between simple walking (SW) and walking with RAS at frequency equal to the subject-specific average cadence (100% AC). Stride Length; Double Limb Support Time (DLS Time); Trunk Displacement Index (TDI); Coefficient of Variability of Double Limb Support Time (DLS Time CV); Significance p value: * p < 0.05, ** p < 0.01, *** p < 0.001.

### 3.2 Low Frequencies

#### 3.2.1 Simple Walking - 90%AC Frequency

The comparison between SW and the 90% AC frequency showed differences in the Temporal Parameters. In particular, the reduced frequency implied a statistically significant decrease of Speed (p < 0.05) and an increase of Stance Time (p < 0.001), Cycle time (p < 0.001) and DLS time (p < 0.001). However, no significant difference was found in Spatial Parameters and in the TDI value in the comparison between these two conditions (Fig. 3).

**Fig. 3.**
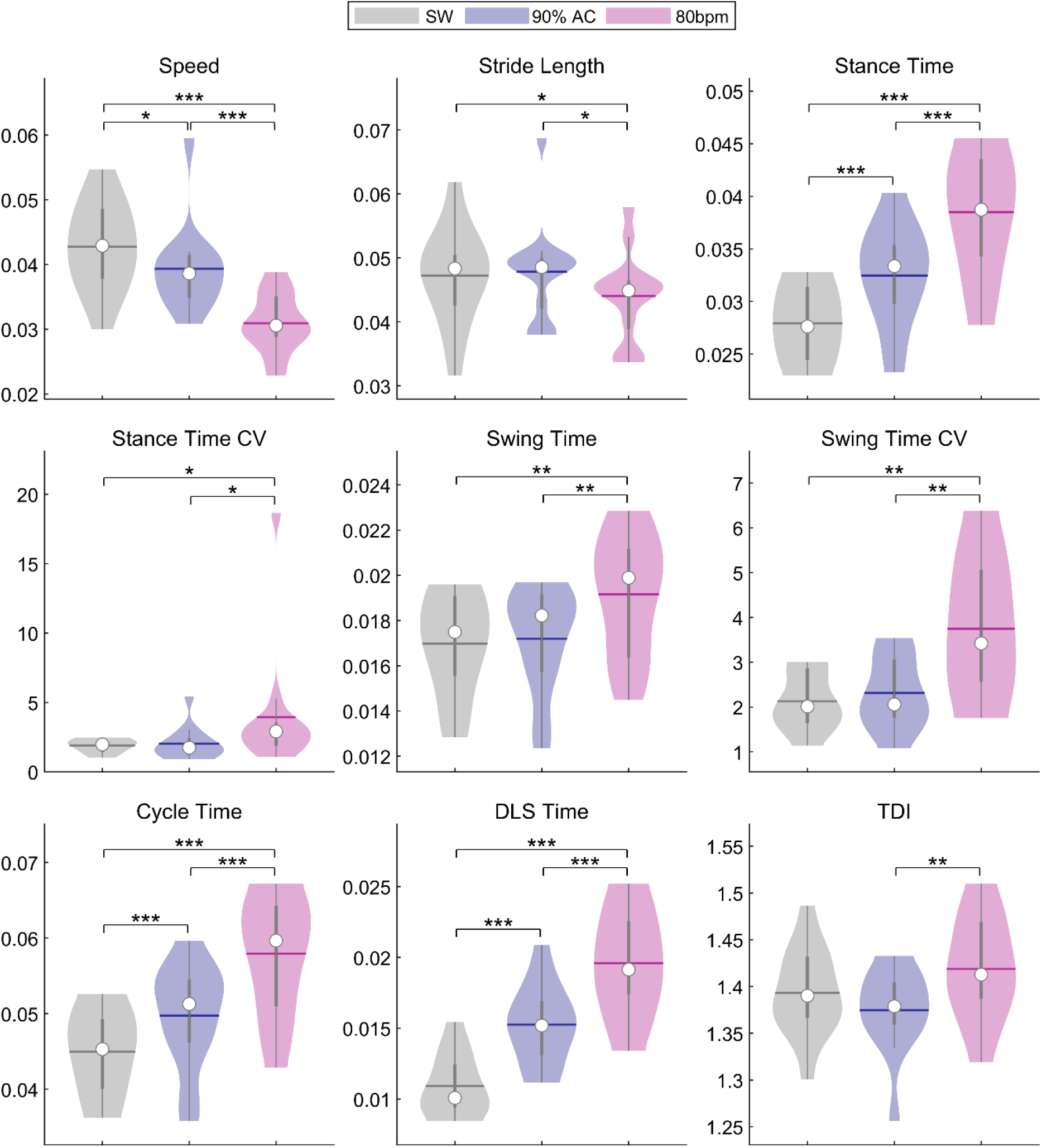
Spatio-temporal analysis of gait and Trunk Displacement Index (TDI). SW vs. Low Frequencies RAS. Violin plots of spatio-temporal parameters and TDI value. Comparison between simple walking (SW) and walking with RAS at low frequencies (SW - 90% AC - 80 bpm). Speed; Stride Length; Stance Time; Coefficient of Variability of Stance time (Stance Time CV); Swing time; Coefficient of Variability of Swing time (Swing Time CV); Cycle Time; Double Limb Support Time (DLS Time); Trunk Displacement Index (TDI); Coefficient of Variability of Double Limb Support Time (DLS Time CV); Significance p value: * p < 0.05, ** p < 0.01, *** p < 0.001.

#### 3.2.2 Simple Walking - 80 bpm Frequency

With regard to the Temporal Parameters, the walking at fixed frequency set at 80 bpm caused a statistically significant reduction of the Speed (p < 0.001), and an increase of the Stance time (p < 0.001), Swing time (p < 0.01), Cycle time (p < 0.001) and DLS time (p < 0.001) as compared to SW. For the Spatial Parameters, it was shown a decrease of the Stride Length (p < 0.05). Furthermore, this frequency of stimulation also caused increased Stance time CV (p < 0.05) and Swing time CV (p < 0.01). No significant difference was found in the TDI between these two conditions (Fig. 3).

### 3.3 High Frequencies

#### 3.3.1 Simple Walking - 110%AC Frequency

The results showed that the 110% AC frequency affected all Temporal Parameters. In particular, it was documented statistically significant increase of Speed (p < 0.001) and decrease of Stance time (p < 0.05), Swing Time (p < 0.001) and Cycle time (p < 0.001), compared to the SW. Concerning the Spatial Parameters, this frequency of stimulation caused a rise of Stride Length (p < 0.05). Moreover, the results showed a reduction of the TDI compared to SW (p < 0.01) (Fig. 4).

**Fig. 4.**
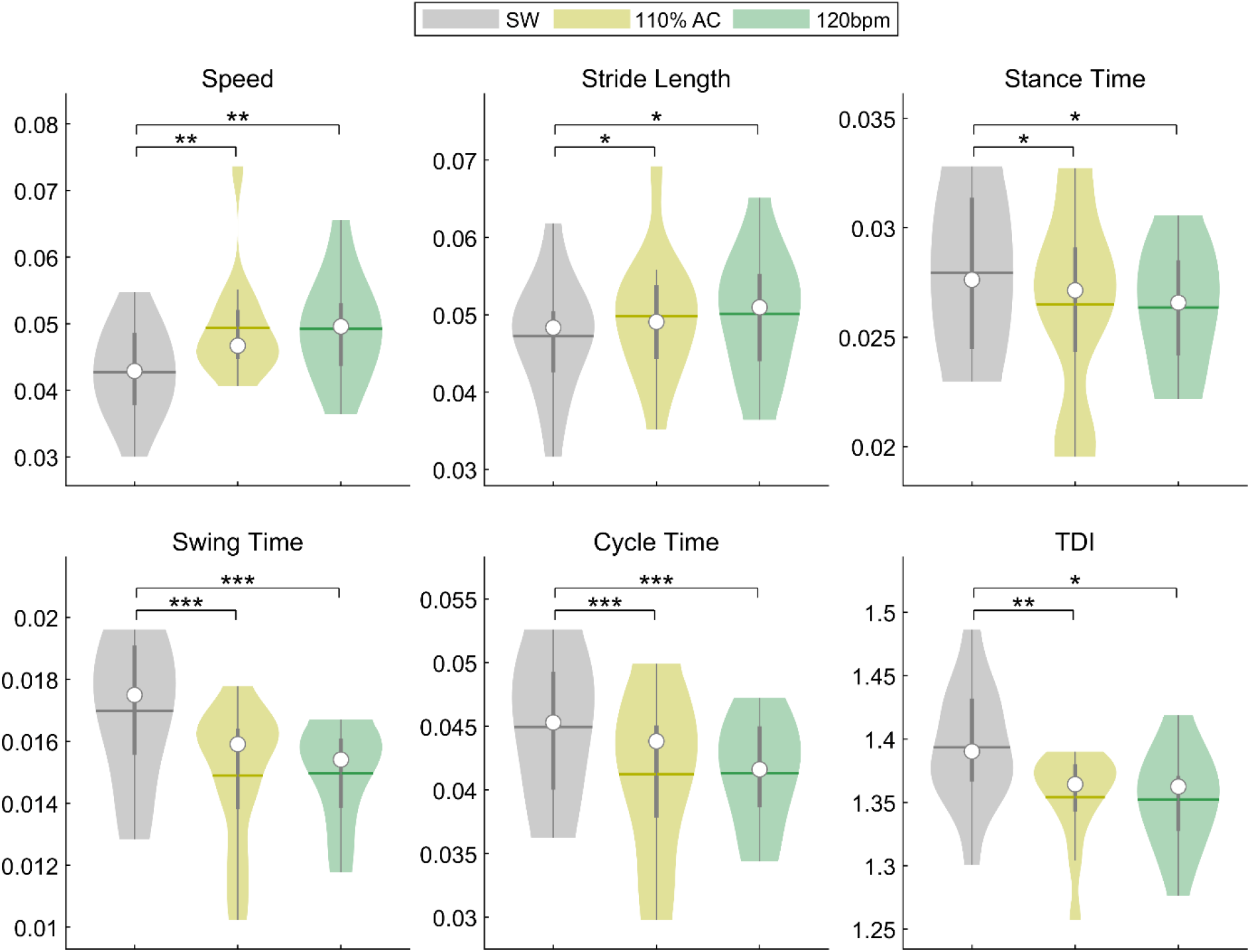
Spatio-temporal analysis of gait and Trunk Displacement Index (TDI). SW vs. High Frequencies RAS. Violin plots of spatio-temporal parameters and TDI value. Comparison between simple walking (SW) and walking with RAS at high frequencies (SW - 110% AC – 120bpm). Speed; Stride Length; Stance Time; Swing time; Cycle Time; Trunk Displacement Index (TDI); Coefficient of Variability of Double Limb Support Time (DLS Time CV); Significance p value: * p < 0.05, ** p < 0.01, *** p < 0.001.

#### 3.3.2 Simple Walking - 120 bpm Frequency

The comparison between the SW and walking at the RAS at 120 bpm showed results resembling that at the 110% AC frequency. In fact, the results highlighted a rise of the Speed (p < 0.001), and a reduction of the Stance time (p < 0.01), Swing time and Cycle time (p < 0.001) with the RAS at 120 bpm. Moreover, our data showed increased Stride length (p < 0.05) and reduced TDI (p < 0.05) (Fig. 4).

### 3.4 Correlations

We performed a correlation analysis to explore the relationship between the TDI and the gait parameters. The TDI correlated negatively with Speed (r = -0.69, p < 0.001) and Stride Length (r = - 0.56, p < 0.001), and positively with Stance Time (r = 0.32, p = 0.012), Swing Time CV (r = 0.27, p = 0.033), Cycle Time (r = 0.32, p = 0.013), and DLS time (r = 0.39, p = 0.002) (Figure 5).

**Fig. 5.**
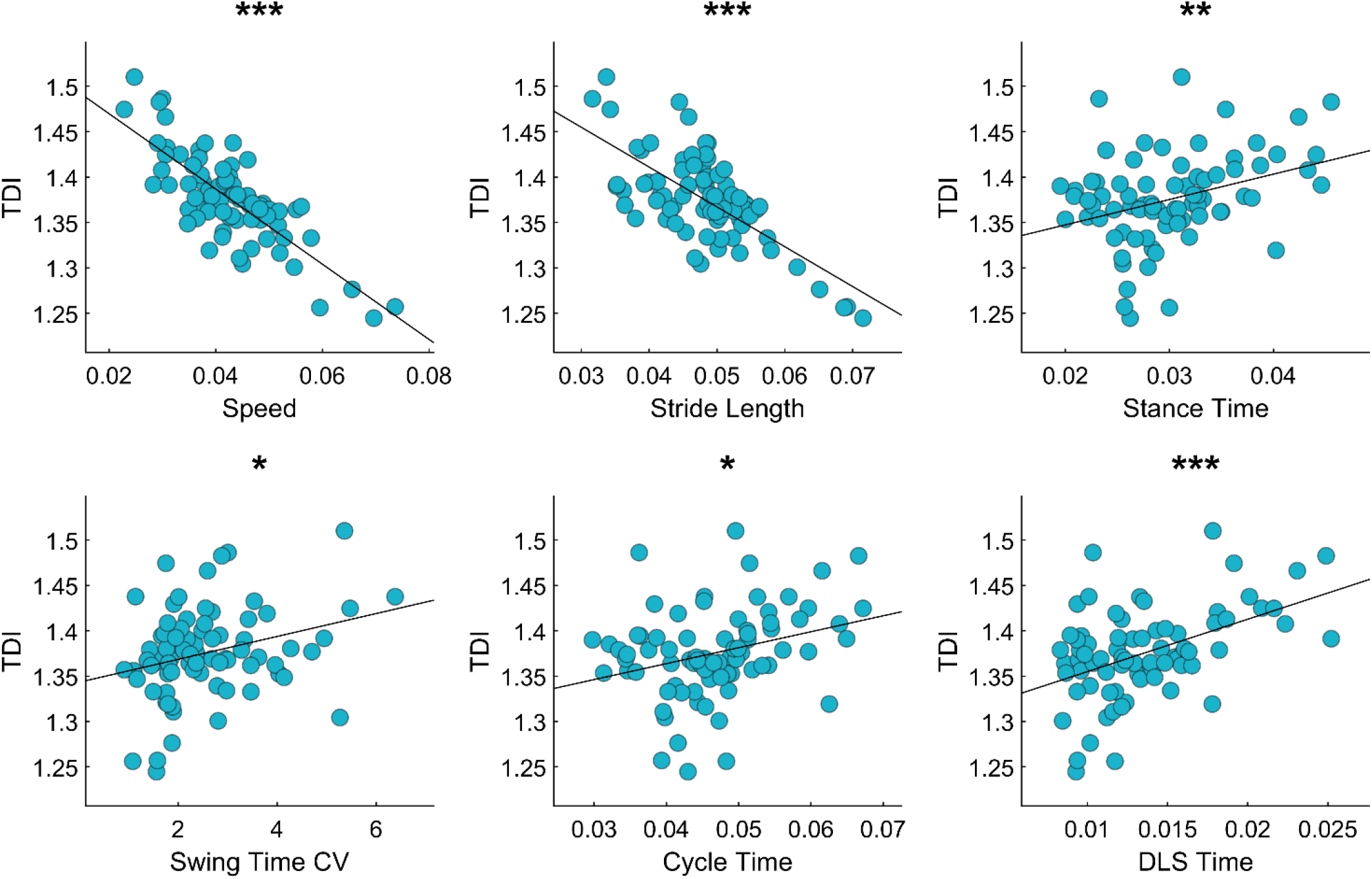
TDI and spatio-temporal gait parameters. Pearson coefficient correlation between trunk displacement index (TDI) and spatio-temporal gait parameters in all conditions. Speed, Stride length, Stance time, Coefficient of variability of Swing Time (Swing Time CV), Cycle Time. Significance p value: * p < 0.05, ** p < 0.01, *** p < 0.001.

## 4. DISCUSSION

In this study, we assessed how different frequencies of the RAS may affect walking in healthy elderly subjects, evaluating the spatio-temporal parameters, and the TDI.

### 4.1 Spatio-Temporal Parameters

#### 4.1.1 SW and 100% AC

As expected, with regard to the comparison between SW and 100% AC, the results highlighted that, by setting the frequency of the RAS equal to the cadence of each subjects, the temporal parameters (speed, stance time, swing time and cycle time) showed no significant change, except for the DLS time, whereas differences in the spatial parameters (Stride length and DLS CV) were observed. In fact, while the results of the temporal parameters showed a synchronization with the RAS, the spatial parameters demonstrated an increase in stability, as suggested by the increase of Stride length and the reduction DLS CV. Data also revealed a raise of DLS time, commonly related to altered stability and increased risk of fall^44–46^. We assume that the DLS time increase is due to a mechanism of synchronization with the stimulus, which involves a longer expectation of the stimulus with both feet in contact with the ground. To overcome this limitation, we calculated the TDI, a measure capturing gait stability. Taking into consideration the trunk displacement in relation to the COM of the individual, the TDI provides a proxy of stability of the whole-body. Indeed, during the 100% AC frequency stimulation, the TDI value decreased, showing a reduction of trunk swings, and therefore a greater stability. This agrees with a study performed by Arias et al. on the effects of rhythmic sensory stimulation on gait in Parkinsonians patients and age-matched healthy controls. Furthermore, Arias et al. also reported increased speed and reduced stride time variability in the same subjects^47^. This partial discrepancy could be caused by the difference in age of the population. Indeed, our participants synchronised with the stimulus and increased the DLS time, rather than the speed, obtaining a reduction of the DLS CV.

#### 4.1.2 Low Frequencies

The results of the comparison between SW and low frequencies stimulation (90% AC and 80 bpm) showed the same trend for almost all parameters, with the main variations in the temporal parameters. In fact, for both stimulations, a reduction of Speed, and a rise of Stance time, Cycle Time and DLS time was observed. In fact, similarly of the 100% AC frequency, even in the low frequencies the increase of the DLS time could be caused by the same mechanism of synchronization with the stimulus. We speculated that, in order to adapt to the low frequency stimulus, the participants increased the whole cycle time by increasing the time spent in double stance, in order to wait for the next stimulus in a condition of higher stability (both feet on the ground, instead of one, as in the swing phase). The gait modification induced by the RAS at 90% frequency in our sample are in agreement with a study performed by Willems et al., which found reduced speed and cadence and unmodified stride length^48^. When reducing the frequency of the stimulus, it can be observed that all the temporal parameters slowed down. On top of that, the lowest frequency (80 bpm) also showed increased Swing time and rise of Stance Time CV and Swing Time CV. To help interpretation, the TDI variations of the two stimulations showed opposite trends. In fact, during the 90% AC frequency stimulation (i.e. the one that was more similar to the natural cadence) we observed a reduction of the TDI value, and hence greater stability. This shows that the differences in the spatio-temporal parameters induced by the stimulus, reduced oscillations and results in enhanced stability, by forcing the subject to walk slightly slower. However, during the 80-bpm frequency stimulation the stability becomes sub-optimal (i.e. forcing the subject to walk at a much lower cadence as shown in table 2. Intuitively, this phenomenon could be thought as similar to the common experience of riding a bicycle: a slow pace makes the riding easier, until the point when the lack of speed makes oscillations more pronounced, and stability harder to maintain. In other words, the modification of the spatio-temporal parameters succeeds at maintaining optimal balance, but fail when the difference becomes too big. This has implication when designing balance-enhancing strategies such as with physiotherapy.

#### 4.1.3 High Frequencies

Also, in this case, the results of the comparison between SW and high frequencies (110% AC and 120 bpm) showed the same trend for all the parameters. In fact, for the Spatial Parameters, one can observe longer Stride Length and, for Temporal Parameters, increased Speed, with the consequent reduction of the Stance Time, the Swing Time and the Cycle Time. Our findings are in accordance with Yu et al., which found increased Speed, cadence and Stride Length in healthy young subjects^30^. It can be observed that the high frequencies do not cause alterations in the double support time. Moreover, both frequencies caused lower TDI (hence smaller oscillations of the trunk over the COM). Therefore, the gait changes that occur under the higher RAS suggest a better ability to control the body sway and consequently less body instability.

### 4.3 Correlations

Our analysis showed that the TDI correlated positively with the DLS time. This finding implies that, when the TDI increases, the DLS increases too. This seems logical as double stance time is the most stable part of the gait cycle^49^, and it typically becomes longer when more stability is requested. Hence, when instability is present, as indicated by higher TDI, the gait is modified toward higher double support time. In fact, the relationship between the TDI and the DLS time is not the same in all conditions. While all the three conditions (100% AC, 90% AC, 80 bpm) saw an increase in DLS time, only the 80 bpm condition showed increased TDI, while the opposite happened in the 100% AC and 90% AC conditions. Our interpretation of this finding is that the subjects successfully responded to the 100% AC RAS and the 90% AC RAS by adapting the DLS phase, in such a way as to obtain more stability, perhaps taking advantage of the external cueing and, consequently, decreasing the oscillations of the trunk. However, the 80 bpm RAS is much lower than the natural walking rhythm of our population, and the over-extended duration of the DLS provoked a deficit in stability, resulting in higher TDI values. In other words, the TDI grows linearly with the DLS for a certain range of stimulation. However, out of this range, when the stimulus becomes too different form the natural frequency, such relationship no longer holds, and higher DLS starts to correspond to lower stability, perhaps as if the compensatory mechanisms can no longer achieve stability. In the 110% AC and 120 bpm cases, we did not observe any significant change in the DLS time, while the TDI decreased significantly. This information suggests that the rise of the TDI might be interpreted as an indicator of a condition of instability.

Moreover, the TDI resulted to correlate negatively with the Speed. Consequently, we can observe that Speed values increase when the TDI values decrease. Significant increase and decrease of Speed were showed during high and low frequency RAS, respectively. As shown by our results, the participants adjusted the Cycle Time, in order to follow the low or high RAS, respectively. Hence, the strategy put in place by participants, to reach the correct Cycle Time, entailed modulating the Speed. Consequently, walking faster caused shorter duration of the Cycle Time. Again, the example of the bicycle can be of help, given that a slightly higher speed can stabilize oscillations, similarly to what we observed in this case.

The TDI was correlated negatively to the Stride Length too. Similarly, to our previous line of thinking, also in this case the presence of optimal balance, as signalled by the low TDI, would mean that it is safer to make longer steps.

Our results agree with several studies which reported that the elderly commonly showed a reduced speed and short stride length, linked to an impaired stability. In a review on balance and gait in the elderly, Osoba et al., describe the slow speed and the short stride length as a cautious gait pattern put in place to increase stability^50^. Furthermore, a clinical guide presented by Pirker and Katzenschlager on gait disorder in elderly people, states that ageing is related to a reduction in step length and, consequently, in speed, highlighting the relationship between the speed and the general health of the subjects^51^.

The positive correlation between TDI and Swing Time CV suggested that the trunk displacement changes are directly proportional to the variability of gait. We cannot prove the directionality of the implication, but one can speculate according to the following line of reasoning. Since the balance control employed by the cerebellum acts on the COM, gathering vestibule-spinal information of trunk verticality^32,35^, it could be hypothesized that an impaired balance control of the upper body, as captured by increased TDI, which contains information of both trunk and COM, could determine instability during both the stance and swing phases of gait. Again, Osoba et al., reported increased gait variability associated with ageing and risk of fall^50^.

## 5. CONCLUSIONS

Our study shows that different RAS can influence the gait parameters in a frequency-specific manner, which means that a frequency equal to the individual natural frequency improve the stability. While low frequencies shows a general slowing down of gait, which do not provide any clear beneficial effect in terms of stability, an excessive low stimulation (80 bpm) seems detrimental to stability, being too slow compared to the individual natural frequency. In accordance with our expectations, using a frequency that exceeds a threshold value causes a worsening of the gait parameters and stability.

The high frequencies of RAS however provokes a slight speeding up of gait, which is accompanied by improved stability. Importantly, our study shows that a moderate increase of Speed is important to positively influence stability. In this case, fixed and variable RAS shared similar frequencies, hence no difference was found in gait with 110% AC and 120 bpm. This result supports our hypothesis that the effectiveness of the stimulation on stability depends on how far the stimulus is from the individual average cadence.

Eventually, the increased stability offered by the RAS may be used both in rehabilitation protocols and prevention strategies. RAS could support classical rehabilitation, adding a sensory training to the purely motor one, tailoring the quality and quantity of the stimulation on each individual. Moreover, the prevention techniques could be aided by the use of devices based on the RAS in order to support the gait continuously, increasing the stability of the individual and preventing them from falls.

One of the limitations of this study is the small data sample, due to the difficulty in finding and recruiting cognitively and physically healthy elderly subjects. Another one, is that the two high frequencies were quite similar to each other’s, hence we could not explore the effects of a stimulation with a frequency much higher than the average cadence of each subject, as for the low frequencies. Future researches on the effects of RAS on the gait could aim to identify the threshold values above which there is a worsening of the gait and a reduction in stability.

## ACKNOWLEDGEMENT

We would like to thank the Centro Sociale Polifunzionale “Mariano Bottari”, Portici (Naples) in the recruitment the subjects.

## BIBLIOGRAPHY

1. Borzuola R, Giombini A, Torre G, et al. Central and Peripheral Neuromuscular Adaptations to Ageing. J Clin Med. 2020;9(3):741.

2. Buckles VD. Age-related slowing. In: Sensorimotor Impairment in the Elderly. Springer; 1993:73–87.

3. Tang P-F, Woollacott MH. Balance control in older adults: Training effects on balance control and the integration of balance control into walking. In: Advances in Psychology. Vol 114. Elsevier; 1996:339–367.

4. Contreras-Vidal JL, Teulings H, Stelmach G. Elderly subjects are impaired in spatial coordination in fine motor control. Acta Psychol (Amst). 1998;100(1-2):25–35.

5. Rucco R, Sorriso A, Liparoti M, et al. Type and location of wearable sensors for monitoring falls during static and dynamic tasks in healthy elderly: a review. Sensors. 2018;18(5):1613.

6. Burns E, Kakara R. Deaths from falls among persons aged≥ 65 years—United States, 2007– 2016. Morb Mortal Wkly Rep. 2018;67(18):509.

7. Coutinho ESF, Bloch KV, Coeli CM. One-year mortality among elderly people after hospitalization due to fall-related fractures: comparison with a control group of matched elderly. Cad Saude Publica. 2012;28:801–805.

8. Padrón-Monedero A, Damián J, Martin MP, Fernández-Cuenca R. Mortality trends for accidental falls in older people in Spain, 2000-2015. BMC Geriatr. 2017;17(1):1–7.

9. Bernard Demanze Laurence LM. THE FALL IN OLDER ADULTS: PHYSICAL AND COGNITIVE PROBLEMS Send Orders for Reprints to The Fall in Older Adults: Physical and Cognitive Problems. Curr Aging Sci. 2017;10(June):185–200. doi:10.2174/1874609809666160630

10. Auvinet B, Touzard C, Montestruc F, Delafond A, Goeb V. Gait disorders in the elderly and dual task gait analysis: a new approach for identifying motor phenotypes. J Neuroeng Rehabil. 2017;14(1):7.

11. Kaczmarczyk K, Wiszomirska I, Błażkiewicz M, Wychowański M, Wit A. First signs of elderly gait for women. Med Pr. 2017;68(4):441.

12. Virmani T, Gupta H, Shah J, Larson-Prior L. Objective measures of gait and balance in healthy non-falling adults as a function of age. Gait Posture. 2018;65(July 2017):100–105. doi:10.1016/j.gaitpost.2018.07.167

13. Dugan SA, Bhat KP. Biomechanics and analysis of running gait. Phys Med Rehabil Clin N Am. 2005;16(3):603–621. doi:10.1016/j.pmr.2005.02.007

14. Boyer KA, Johnson RT, Banks JJ, Jewell C, Hafer JF. Systematic review and meta-analysis of gait mechanics in young and older adults. Exp Gerontol. 2017;95(2016):63–70. doi:10.1016/j.exger.2017.05.005

15. Liparoti M, Della Corte M, Rucco R, et al. Gait abnormalities in minimally disabled people with Multiple Sclerosis: A 3D-motion analysis study. Mult Scler Relat Disord. 2019;29. doi:10.1016/j.msard.2019.01.028

16. Sorrentino P, Barbato A, Del Gaudio L, et al. Impaired gait kinematics in type 1 Gaucher’s Disease. J Parkinsons Dis. 2016;6(1):191–195. doi:10.3233/JPD-150660

17. Pistacchi M, Gioulis M, Sanson F, et al. Gait analysis and clinical correlations in early Parkinson’s disease. Funct Neurol. 2017;32(1):28–34. doi:10.11138/FNeur/2017.32.1.028

18. Amboni M, Iuppariello L, Iavarone A, et al. Step length predicts executive dysfunction in Parkinson’s disease: a 3-year prospective study. J Neurol. 2018;265(10):2211–2220.

19. Rucco R, Liparoti M, Agosti V. A new technical method to analyse the kinematics of the human movements and sports gesture. J Phys Educ Sport. 2020;20(4):2360–2363. doi:10.7752/jpes.2020.s4319

20. Petersen H, Magnusson M, Johansson R, Akesson M, Fransson PA. Acoustic cues and postural control. Scand J Rehabil Med. 1995;27(2):99–104.

21. Kristinsdottir M. Magnusson, EK PAF. Changes in postural control in healthy elderly subjects are related to vibration sensation, vision and vestibular asymmetry. Acta Otolaryngol. 2001;121(6):700–706.

22. Wittwer JE, Webster KE, Hill K. Music and metronome cues produce different effects on gait spatiotemporal measures but not gait variability in healthy older adults. Gait Posture. 2013;37(2):219–222. doi:10.1016/j.gaitpost.2012.07.006

23. Rhea CK, Kuznetsov NA. Using visual stimuli to enhance gait control. J Vestib Res Equilib Orientat. 2017;27(1):7–16. doi:10.3233/VES-170602

24. Dozza M, Horak FB, Chiari L. Auditory biofeedback substitutes for loss of sensory information in maintaining stance. Exp Brain Res. 2007;178(1):37–48. doi:10.1007/s00221-006-0709-y

25. Thaut M, McIntosh G, Rice R. Rhythmic auditory stimulation as an entrainment and therapy technique in gait of stroke and Parkinson’s patients. Music II. 1995.

26. Howe T, Lovgreen B, Cody F.V. Ashtone J. Oldham,”Auditory cues can modify the gait of persons with early-stage Parkinson’s disease: a method for enhancing parkinsonian walking performance?,.” Clin Rehabil. 2003;17(4):363–367.

27. Bella SD, Benoit CE, Farrugia N, et al. Gait improvement via rhythmic stimulation in Parkinson’s disease is linked to rhythmic skills. Sci Rep. 2017;7(February):1–11. doi:10.1038/srep42005

28. Shahraki M. Effect of rhythmic auditory stimulation on gait in patients with stroke. Parkinsonism Relat Disord. 2016;22(1):e125. doi:10.1016/j.parkreldis.2015.10.295

29. Dickstein R, Plax M. Metronome Rate and Walking Foot Contact Time in Young Adults. Percept Mot Skills. 2012;114(1):21–28. doi:10.2466/15.25.PMS.114.1.21-28

30. Yu L, Zhang Q, Hu C, Huang Q, Ye M, Li D. Effects of different frequencies of rhythmic auditory cueing on the stride length, cadence, and gait speed in healthy young females. J Phys Ther Sci. 2015;27(2):485–487. doi:10.1589/jpts.27.485

31. van Mastrigt NM, Celie K, Mieremet AL, Ruifrok ACC, Geradts Z. Critical review of the use and scientific basis of forensic gait analysis. Forensic Sci Res. 2018;3(3):183–193. doi:10.1080/20961790.2018.1503579

32. Takakusaki K. Functional neuroanatomy for posture and gait control. J Mov Disord. 2017;10(1):1.

33. Grasso R, Zago M, Lacquaniti F. Interactions between posture and locomotion: motor patterns in humans walking with bent posture versus erect posture. J Neurophysiol. 2000;83(1):288–300.

34. Lopez ET, Minino R, Sorrentino P, et al. A synthetic kinematic index of trunk displacement conveying the overall motor condition in Parkinson’s disease. bioRxiv. 2020.

35. Jo S. Hypothetical neural control of human bipedal walking with voluntary modulation. Med Biol Eng Comput. 2008;46(2):179–193.

36. Beck AT, Steer RA, Brown G. Beck depression inventory–II. Psychol Assess. 1996.

37. Folstein MF, Folstein SE, McHugh PR. “Mini-mental state”: a practical method for grading the cognitive state of patients for the clinician. J Psychiatr Res. 1975;12(3):189–198.

38. Dubois B, Slachevsky A, Litvan I, Pillon B. The FAB: A frontal assessment battery at bedside. Neurology. 2000;55(11):1621–1626. doi:10.1212/WNL.55.11.1621

39. Davis RB, Õunpuu S, Tyburski D, Gage JR. A gait analysis data collection and reduction technique. Hum Mov Sci. 1991;10(5):575–587. doi:10.1016/0167-9457(91)90046-Z

40. Vitale C, Agosti V, Avella D, et al. Effect of Global Postural Rehabilitation program on spatiotemporal gait parameters of parkinsonian patients: a three-dimensional motion analysis study. Neurol Sci. 2012;33(6):1337–1343.

41. Coste A, Salesse RN, Gueugnon M, Marin L, Bardy BG. Standing or swaying to the beat: Discrete auditory rhythms entrain stance and promote postural coordination stability. Gait Posture. 2018;59(January 2017):28–34. doi:10.1016/j.gaitpost.2017.09.023

42. Almarwani M, Van Swearingen JM, Perera S, Sparto PJ, Brach JS. The Effect of Auditory Cueing on the Spatial and Temporal Gait Coordination in Healthy Adults. J Mot Behav. 2017;2895:1–7. doi:10.1080/00222895.2017.1411330

43. Benjamini Y, Hochberg Y. Controlling the false discovery rate: a practical and powerful approach to multiple testing. J R Stat Soc Ser B. 1995;57(1):289–300.

44. Scott D, McLaughlin P, Nicholson GC, et al. Changes in gait performance over several years are associated with recurrent falls status in community-dwelling older women at high risk of fracture. Age Ageing. 2015;44(2):287–293.

45. Doyo W, Kozakai R, Kim H, Ando F, Shimokata H. Spatiotemporal components of the 3-D gait analysis of community-dwelling middle-aged and elderly Japanese: Age-and sex-related differences. Geriatr Gerontol Int. 2011;11(1):39–49.

46. Williams DS, Martin AE. Gait modification when decreasing double support percentage. J Biomech. 2019;92:76–83.

47. Arias P, Cudeiro J. Effects of rhythmic sensory stimulation (auditory, visual) on gait in Parkinson’s disease patients. Exp brain Res. 2008;186(4):589–601.

48. Willems AM, Nieuwboer A, Chavret F, et al. The use of rhythmic auditory cues to influence gait in patients with Parkinson’s disease, the differential effect for freezers and non-freezers, an explorative study. Disabil Rehabil. 2006;28(11):721–728. doi:10.1080/09638280500386569

49. Xu R, Wang X, Yang J, et al. Comparison of the COM-FCP inclination angle and other mediolateral stability indicators for turning. Biomed Eng Online. 2017;16(1):1–15.

50. Osoba MY, Rao AK, Agrawal SK, Lalwani AK. Balance and gait in the elderly: A contemporary review. Laryngoscope Investig Otolaryngol. 2019;4(1):143–153.

51. Pirker W, Katzenschlager R. Gait disorders in adults and the elderly. Wien Klin Wochenschr. 2017;129(3-4):81–95

